# Structural analysis of nonapeptides derived from elastin

**DOI:** 10.1101/868257

**Authors:** B Hernandez, JM Crowet, J Thiery, SG Kruglik, N Belloy, S Baud, M Dauchez, L Debelle

## Abstract

Elastin-derived peptides are released from the extracellular matrix remodeling by numerous proteases and seem to regulate many biological processes, notably cancer progression. The canonical elastin peptide is VGVAPG which harbors the XGXXPG consensus pattern allowing interaction with the elastin receptor complex located at the surface of cells. Besides these elastokines, another class of peptides has been identified. This group of bioactive elastin peptides presents the XGXPGXGXG consensus sequence but the reason for their bioactivity remains unexplained. In order to better understand their nature and structure-function relationships, herein we searched the current databases for this nonapeptide motif and observed that the XGXPGXGXG elastin peptides define a specific group of tandemly repeated patterns. Further, we focused on four tandemly repeated human elastin nonapeptides, *i.e.* AGIPGLGVG, VGVPGLGVG, AGVPGLGVG and AGVPGFGAG. These peptides were analysed by means of optical spectroscopies and molecular dynamics. UV-circular dichroism and Raman spectra are consistent with a conformational equilibrium between β-turn, β-strand and random chain secondary elements in aqueous media. This equilibrium was found to be concentration-independent. Quantitative analysis of their conformations suggested that turns corresponded to half of the total population of structural elements while the remaining half was equally distributed between β-strand and unordered chains. These distributions were confirmed by molecular dynamics simulations. Altogether, our data suggest that these peptides harbor a type II β-turn located in their central part. We hypothesize that this structural element could explain their specific bioactivity.

**Statement of Significance:** Elastin fragmentation products, the so-called elastin peptides, may exhibit a bioactivity towards normal and tumor cells. This phenomenon depends on the sequence motif they harbor. While XGXXPG sequences bioactivity is explained by the presence of a type VIII β-turn allowing interaction with the elastin receptor complex, the structural reasons for XGXPGXGXG specific activity remain unexplained. Using data mining, we show that elastin nonapeptides define a specific class of tandemly repeated features. Further, spectroscopic and numerical simulations methods suggest the presence of a type II β-turn in their conformation. This structural element could explain their bioactivity.

## Introduction

Elastin is the extracellular matrix protein responsible for the structural integrity and function of tissues undergoing reversible extensibility or deformability (1). This protein is extremely stable and resistant and undergoes virtually no turnover. Nevertheless, during aging, mechanical stress and elastase activities contribute to the fragmentation of this macropolymer into elastin-derived peptides (EDP) (2).

Elastin is characterized by its elasticity and seems devoid of any other biological activity (1). In contrast, EDP have been shown to regulate numerous biological processes and are thought to be involved in several aging-related pathologies such as atherosclerosis (3, 4) and cancer (5–8). Most biologically active EDP, elastokines, possess a GXXPG consensus sequence adopting a type VIII β-turn structure (9) involved in their binding to the elastin receptor complex (10). The interaction of GXXPG-containing elastokines with the elastin receptor complex and the consequences of their interaction has been considerably documented during the last decades. For a comprehensive review on this topic, the reader is referred to (11).

Besides the classical GXXPG-containing sequences, another class of elastokines has also been reported. In 1988, Long and colleagues reported that nonapeptide sequences from elastin were chemoattractant for fibroblasts (12). In 1989, they further demonstrated that this biological activity could be extended to endothelial cells (13). Strikingly, since then, this peculiar class of EDP has been mostly ignored until 2007, when Maeda and colleagues demonstrated that these peptides could promote macrophages migration via a specific, but unknown, receptor (14). More recently, their biological activity has been further linked to lung carcinoma progression (8) and to promotion of invasion in triple-negative cancer cells via MMP-14 and MMP-2 (15).

The relatively low interest deserved by these nonapeptide EDP can be explained by the fact that considerable advances have been made on both GXXPG-containing EDP and their receptor biology. As a consequence, few groups considered investigating on these peculiar peptides.

The nonapeptide AGVPGLGVG and other EDPs share similar structural features: random coil and β-turns conformations. However, AGVPGLGVG induces angiogenesis, tumor progression, secretion of proteases higher than other EDPs in carcinomas. Moreover, *in vitro*, it behaves like amyloid-like peptides harboring the XGGXG motif by forming cross β-sheets at the supramolecular level (16). Very recently, Brassart and colleagues have shown that the receptor mediating AGVPGLGVG biological effects, and identified as lactose-insensitive, is the Ribosomal Protein SA (17).

The aim of this work is to analyze the structure and conformational behavior of these peptides. Our structural analysis was conducted using sequence analysis, spectroscopic methods and molecular dynamics simulations.

Our results show that elastin nonapeptides define a peculiar class among peptides harboring the consensus X-G-X-P-G-X-G-X-G sequence. In elastin, these peptides are mostly tandemly repeated. Further our data show that they are engaged in a conformational equilibrium between random coil and turn conformations that appear independent of concentration or temperature. Molecular dynamics simulations suggest that the dominant conformation of these peptides is a type II β-turn. The functional significance of this conformation is discussed.

## Material and Methods

### Sequence analysis

#### Sequence retrieval

Sequences harboring the X-G-X-P-G-X-G-X-G consensus motif were searched using Python scripts on UniProtKB (Swiss-Prot and TrEMBL) database Fasta files where spliced variants were ignored. Only exact matches were retrieved and further cleaned informatically and manually so as to ensure that a sequence belonging to a given species (identified by its unique accession number) was present only once in our results. For each sequence hit, the collected data were its accession number, the originating organism, the length of the protein and the position(s) of the nonapeptide sequence(s).

#### Pattern analysis

Nonapeptide exact matches were classified and distributed as a function of the taxonomic hierarchical classification lineage of the source organism of the corresponding parent protein. Hits were either individual hits, *i.e.* only one occurrence was found in the sequence, or multiple ones. In the latter case, we distinguished three possibilities. Multiple hits could be found in different regions of the protein and each occurrence was isolated from the others. Another possibility was that the repetitions could overlap, *i.e.* the end of the first occurrence (-X-G) was the beginning of the following one (X-G-). Finally, we observed that nonapeptide sequences could also be tandemly repeated, one after the other. The tree figure has been made with the Python module ETE3 (18) and the Logo sequences with WebLogo (https://github.com/weblogo/weblogo) (19).

### Optical spectroscopy

#### Molecular compounds

Powder samples of amino acids were purchased from Sigma-Aldrich (Saint Quentin Fallavier, France). Lyophilized samples of nonapeptides were obtained from Genecust (Luxemburg, Luxemburg) as zwitterionic peptides in acetate salts. Their purity was at least 95% as determined by mass spectrometry.

#### Solution samples

Lyophilized powder of amino acids and peptides was dissolved in pure water from a Millipore filtration system (Guyancourt, France). The ionic strength of all peptide samples was increased by adding 150 mM NaCl to stock solutions. Upon dissolution, the pH was between 4 and 4.5. Raman spectra were recorded between 10°C and 80°C. Circular dichroism (CD) spectra were obtained at room temperature between 100 and 200 µM.

#### CD spectroscopy

Room temperature ultraviolet-circular dichroism (UV-CD) spectra were analyzed on a JASCO J-810 spectrophotometer (Lisses) within the 190–300 nm spectral region. A path length of 1 mm and a spectral resolution of 0.2 nm were selected. Each spectrum corresponding to an average of five scans was recorded with a speed of 100 nm per min. The measured ellipticity for each sample, referred to as [*ϕ*]_*obs*_, was further normalized to obtain the so-called mean residue ellipticity, [*ϕ*], by using the expression [*ϕ*] = [*ϕ*]_*obs*_/*ncl* where *n*, *c*, and *l* are the length of the peptide, its molar concentration, and the optical path length, respectively. The normalized ellipticity was expressed in deg cm^2^ dmol^−1^.

#### Raman spectroscopy

Room temperature Stokes Raman spectra were analyzed in bulk samples at right angle on a Jobin-Yvon T64000 spectrometer (Longjumeau, France) at single spectrograph configuration, 1200 grooves/mm holographic grating and a holographic notch filter. Raman data corresponding to 1200 s acquisition time for each spectrum were collected on a liquid nitrogen cooled CCD detection system (Spectrum One, Jobin-Yvon). The effective slit width was set to 5 cm^−1^. Solution samples were excited by the 488 nm line of an Ar^+^ laser (Spectra Physics, Evry, France), with 200 mW power at the sample.

#### Post-record spectroscopic data treatment

Buffer subtraction and smoothing of the observed spectra was performed using the GRAMS/AI Z.00 package (Thermo Galactic, Waltham, MA, USA). Final presentation of Raman spectra was done by means of SigmaPlot package 6.10 (SPSS Inc., Chicago, IL, USA).

### Atomistic molecular dynamics simulations

Ten simulations of 500 ns were performed for the 4 human elastin nonapeptides using amber99SB-ILDN force field (20) and Gromacs 2016.5 software (21). Peptides were built in an extended conformation with PyMOL (22) and solvated with TIP3P water in a 5 nm cubic box. All studied systems were first minimized by steepest descent for 5000 steps. Then, a 1-ns simulation with the peptides under position restraints was run before the production simulations were performed. Periodic boundary conditions were used with a 2 fs time step. The dynamics were carried out under NPT conditions (310 K and 1 bar). The temperature was maintained using the v-rescale method (23) with τT = 0.1 ps, and an isotropic pressure was maintained using the Parrinello–Rahman barostat (24) with a compressibility of 4.5 × 10^−5^ bar^−1^ and τP = 2 ps. Short-range non-bonded interactions were treated with a cutoff of 1.0 nm and long-range interactions were calculated with the particle mesh Ewald method (25) with a grid spacing of 0.16 nm. Bond lengths were maintained with the LINCS algorithm (26) and long range dispersion corrections for energy and pressure were applied. The 3D structures were analyzed with both PyMOL and VMD (27) softwares. The secondary structures were computed with DSSP (28). Clustering was performed using the gromos method (29) with a cut-off of 0.1 nm.

## Results

### Sequence analysis

The retrieval of sequences possessing the X-G-X-P-G-X-G-X-G yielded 96 293 unique sequences. These sequences (Table S1) were classed as a function of their taxonomy (Figure 1). Most sequences were from *Bacteria* (71 504 sequences for 75 718 occurrences with 2129 overlaps and 297 tandem repeats) then *Eukariota* (22 919 sequences for 24 828 occurrences with 811 overlaps and 297 tandem repeats), *Archae* (1 630 sequences for 1 694 occurrences with 16 overlaps and 6 tandem repeats) and *Virus* (240 sequences for 245 occurrences with no overlaps and no tandem repeats). Strikingly, when these results are observed, it comes that elastins define a specific group. They exhibit numerous occurrences per sequences (171 for 38) with no overlap and a very high ratio of tandem repeats. This specificity of elastin nonapeptides is even more evident when the most frequent residues occurring at positions X are considered. While other group mostly favor the presence of G, A, K, P or L at these positions, elastins allow almost none of those with the AGVPGFGVG being the dominating motif. In fact, out of the 140 million sequences searched, this motif was always found in elastin sequences but one bacterial sequence.

**Figure 1.**
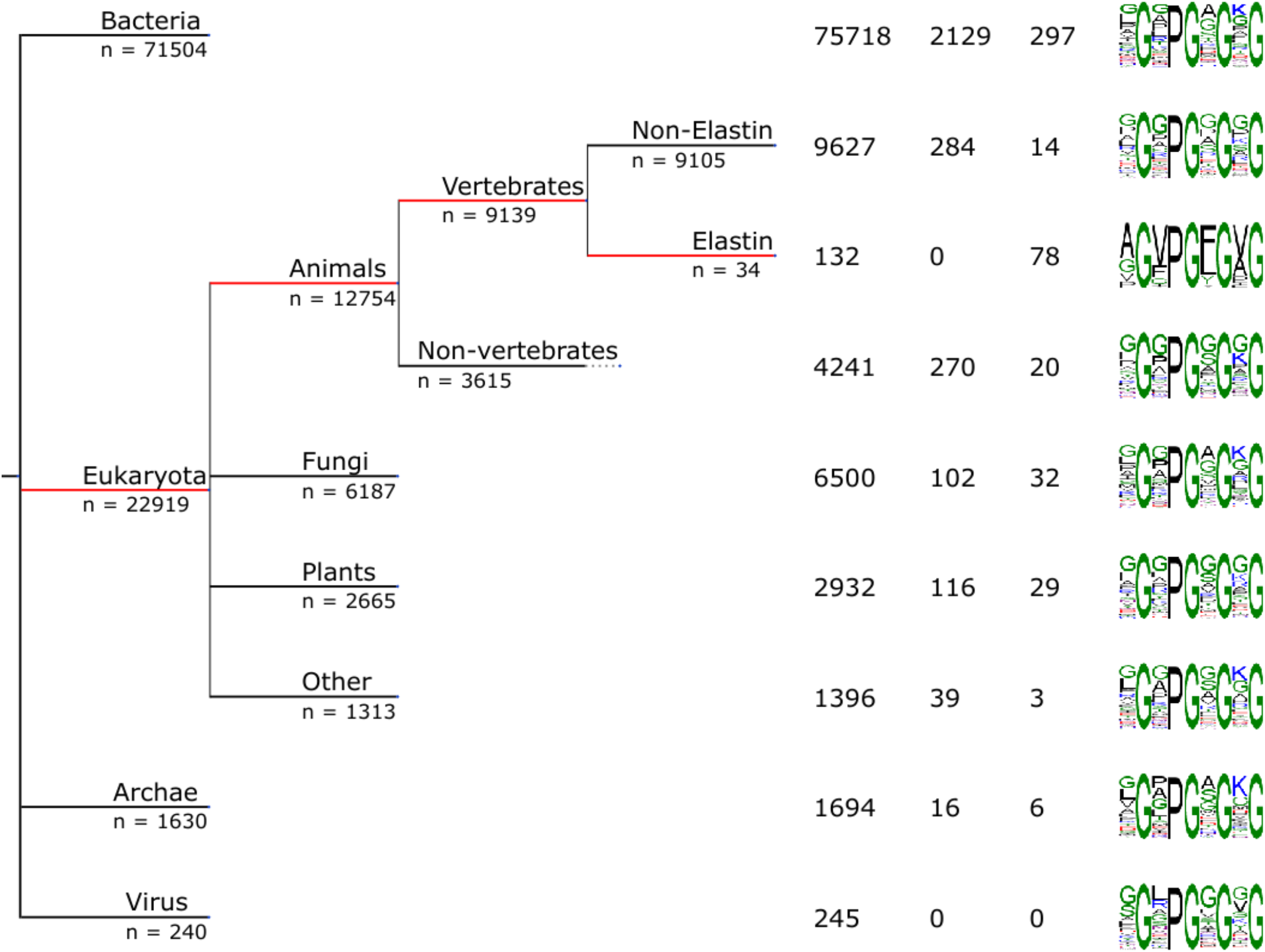
Nonapeptide occurrences and distribution. The X-G-X-P-G-X-G-X-G consensus was found in 96 293 unique sequences amongst which 22 919 were from *Eukariota*, 71 504 from *Bacteria*, 240 from *Virus* and 1630 from *Archae*. The *Eukariota* branch distribution is dominated by Animals in which Vertebrates are predominant. For each branch of the tree, the total number of hits is reported as well as the number of observed overlapping sequences and of tandem repetitions. The consensus motif is also presented with the most frequent residues found at the four X positions.

The nonapeptide sequences found in elastin entries are reported in Table 1 where they are compared. The sequences are reported as a function of their alphabetical order. In human elastin, nonapeptides can be observed in exon domains 18, 20 or 26. Single occurrences are observed in the two first exons while they appear almost exclusively as tandem repeats in exon 26. Strikingly, the human repeats can also be observed in other species. For instance, the human elastin motif AGVPGFGVG is also present in mouse exon 26. Overall, human exon 26 (or its equivalent in other species) is a sequence domain characterized by tandem repeats of these nonapeptides. As a consequence, those are supposedly important for the biological function of the polymer and we decided to further analyze the structural behavior of the 4 human exon 26 nonapeptides, *i.e.* AGIPGLGVG (hN3), VGVPGLGVG (hN4), AGVPGLGVG (hN5) and AGVPGFGAG (hN6).

**Table 1.**
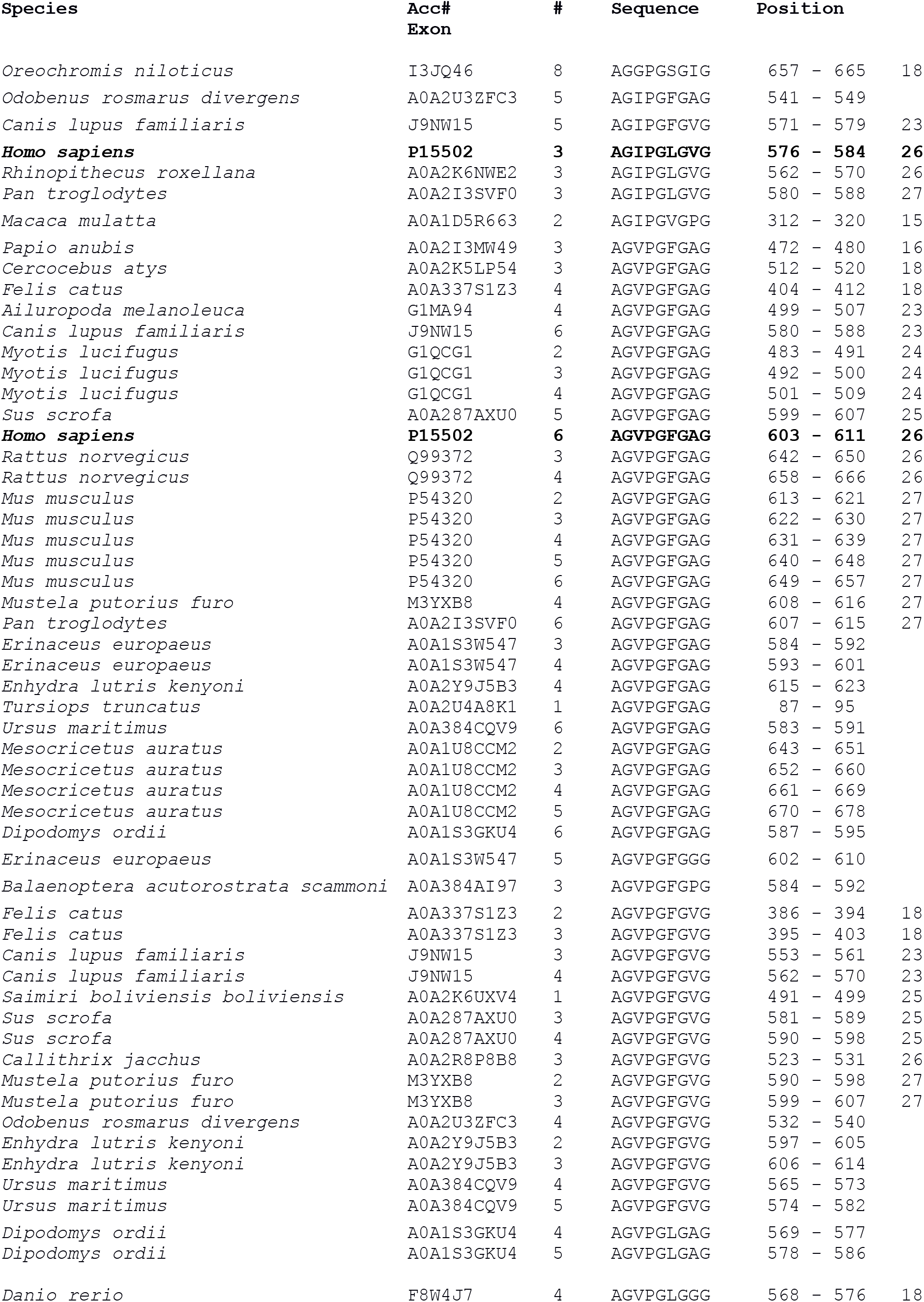
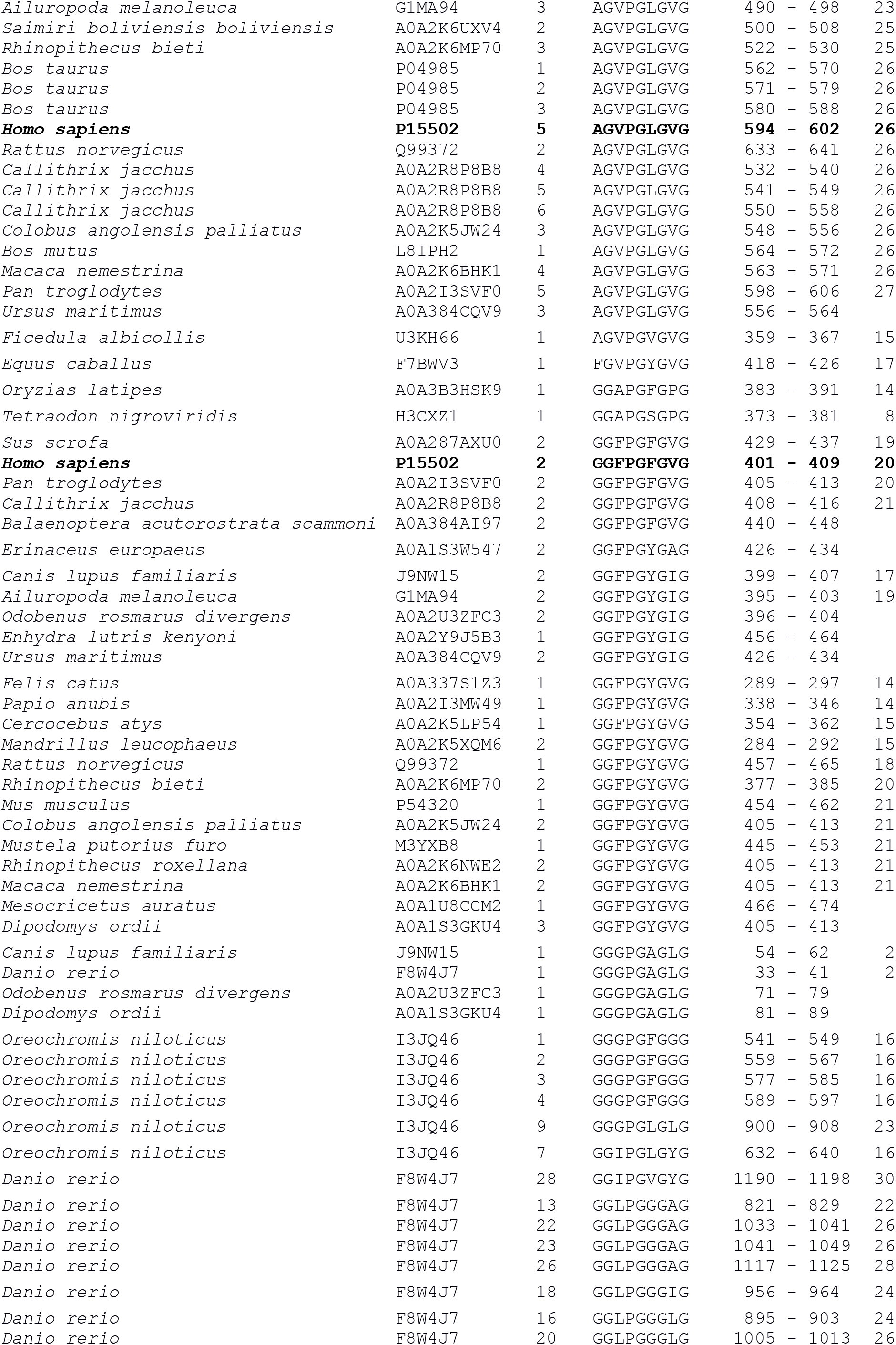
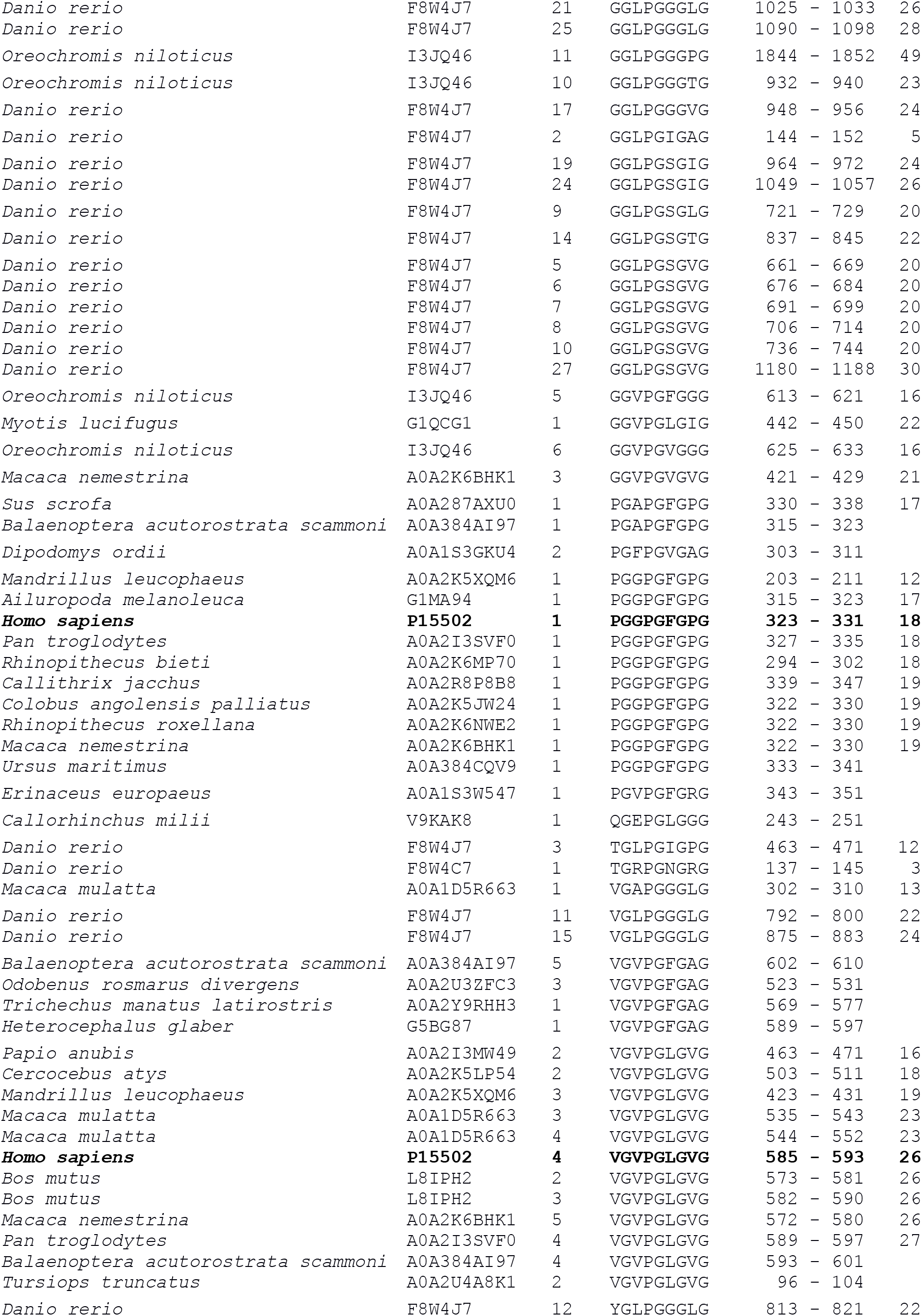
Nonapeptide stack of elastin Uniprot entries Species, name of the species. Acc#, accession number of the origin sequence. #, number of hits in the sequence. Position, position of the hit. Exon, exon number (when available). Human sequences appear in bold faces.

### Structural behavior in the sub-millimolar range

Beyond the simple exploration of the concentration-dependence of the conformational features in aqueous media, our aim was to search the structural behavior of the nonapeptides in different (polar *vs* low dielectric constant) environments. Routinely, methanol is considered as a primary step in studying the peptides conformational behavior in hydrophobic environments, such as membrane interior or hydrophobic pockets of proteins (30).

The analysis of the CD spectra in water (150 mM NaCl), in a water/methanol mixture (50/50, v/v), as well as in methanol (Fig. 2) reveal a progressive conformational change of the nonapeptides in going from water to methanol solutions.

**Figure 2.**
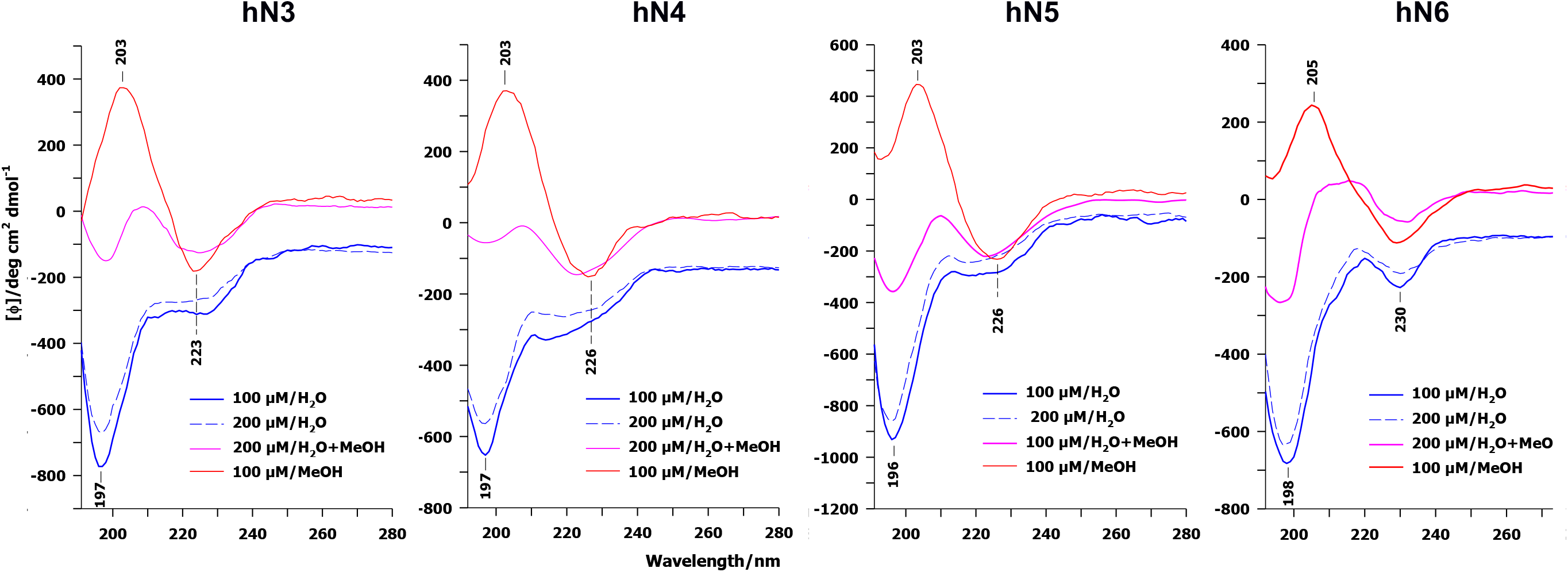
Ultraviolet-circular dichroism (UV-CD) spectra of the four nonapeptides observed in water, water-methanol mixture (50/50), and methanol.

The CD spectra recorded in water all present a double negative band shape, *i.e.* composed of a deep minimum at ~198 nm followed by a weaker and broader negative band centered at ~225 nm. It is a matter of fact that random chains are generally identified by a deep negative band at ~198 nm (31, 32). However, a negative double band, as that observed in the case of the nonapeptides (Fig. 2), is an indicator of turns. For instance, certain β-turns are known to give rise to a negative double band. It is to be mentioned that β-turns are recognized as the most frequent ones in proteins (33)(34)(35). Particularly, the CD fingerprints of four categories of β-turns, i.e. type-I (31), −II and −II’ (36–38) and −VIII (39) β-turns are identified by a double negative band, with unequal ellipticities; the lower wavelength minimum is generally less intense than the higher wavelength one. It is interesting to note that amongst all the mentioned β-turns, that corresponding to type-VIII is formed by a deep minimum at ~198 nm followed by a broad shoulder at ~220 nm (39), *i.e.* strikingly similar to that observed in the case of the presently analyzed nonapeptides. However, this overwhelming resemblance between CD markers does not permit rejection of possible unordered chains, of which the CD marker can be overlapped with the negative band at ~198 nm. In the same manner, a probable overlap may occur between the CD fingerprint of β-strands, generally characterized by a unique negative band at ~215 nm, and the ~225 nm negative band from β-turns. Nevertheless, any uncertainty about the presence of PPII (polyproline II) type chains can be discarded because of the recent CD studies highlighting the fact that the PPII fingerprint is composed of a deep negative band at ~198 nm followed by a positive one centered at ~220 nm. The presence of the latter band cannot be suspected in the CD spectra displayed in Figure 2. At last, the CD spectra of the four nonapeptides recorded in water were found to be concentration-independent in the 100-200 µM range.

In methanol (Fig. 2), the CD signals of the nonapeptides are composed of a broad positive band peaking at ~205 nm followed by a weak and broad negative band at ~225 nm. It is worth noting that the CD spectra observed in water/methanol mixture (1:1, v/v) are somehow an overlap of those observed in water and methanol. To provide a reliable assignment for the CD signal observed in methanol, we can recall that while a type-II β-turn gives rise to a positive band located between 190 and 210 nm (31), the observed negative band at ~225 nm in methanol can be ascribed to β-strands and/or to type-I’ β-turn (31). Based on the observed CD spectra, no unordered chain can be expected in the presently studied nonapeptides.

### Structural behavior in the millimolar range

The CD spectra described in the preceding section pointed out the possible coexistence of several secondary structural elements, such as β-turn, β-strand and unordered chain in the sub-millimolar concentration range. In this section, we checked by means of Raman spectroscopy whether (i) this conformational equilibrium remains concentration-independent upon increasing concentration; (ii) the respective populations of different conformational elements are subjected to a measurable change upon increasing concentration. It should be reminded that, in a general manner, the favorable interactions occurring between the residues having branched aliphatic side chains (such as those existing in the nonapeptides), also named hydrophobic interactions, reinforce peptide aggregation and lead to a subsequent change in the conformational equilibrium within the millimolar concentration range (37).

Raman spectra recorded in the middle wavenumber region (1800-550 cm^−1^) present a low water contribution enabling an accurate analysis of the observed bands (Fig. 3A-B). Whatever the nonapeptide, the Raman spectra were found to be concentration-independent in the 2.5-20 mM interval (data not shown). Indeed, no substantial change assignable to either a possible conformational transition or a detectable molecular aggregation, could been observed. This is the reason why, in the following, we will limit our discussion to the Raman spectra recorded at the ultimate concentration, *i.e.* 20 mM.

**Figure 3.**
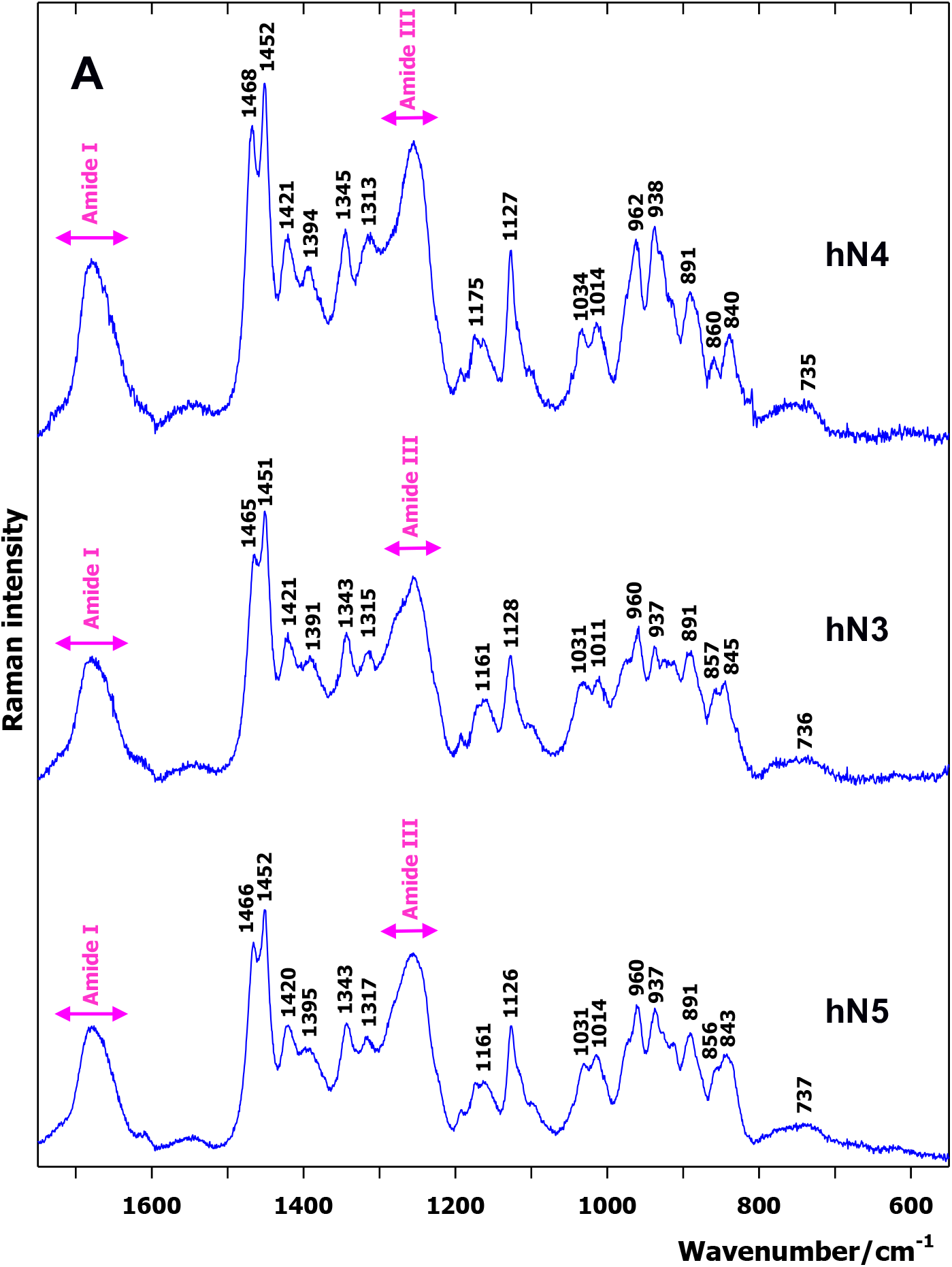
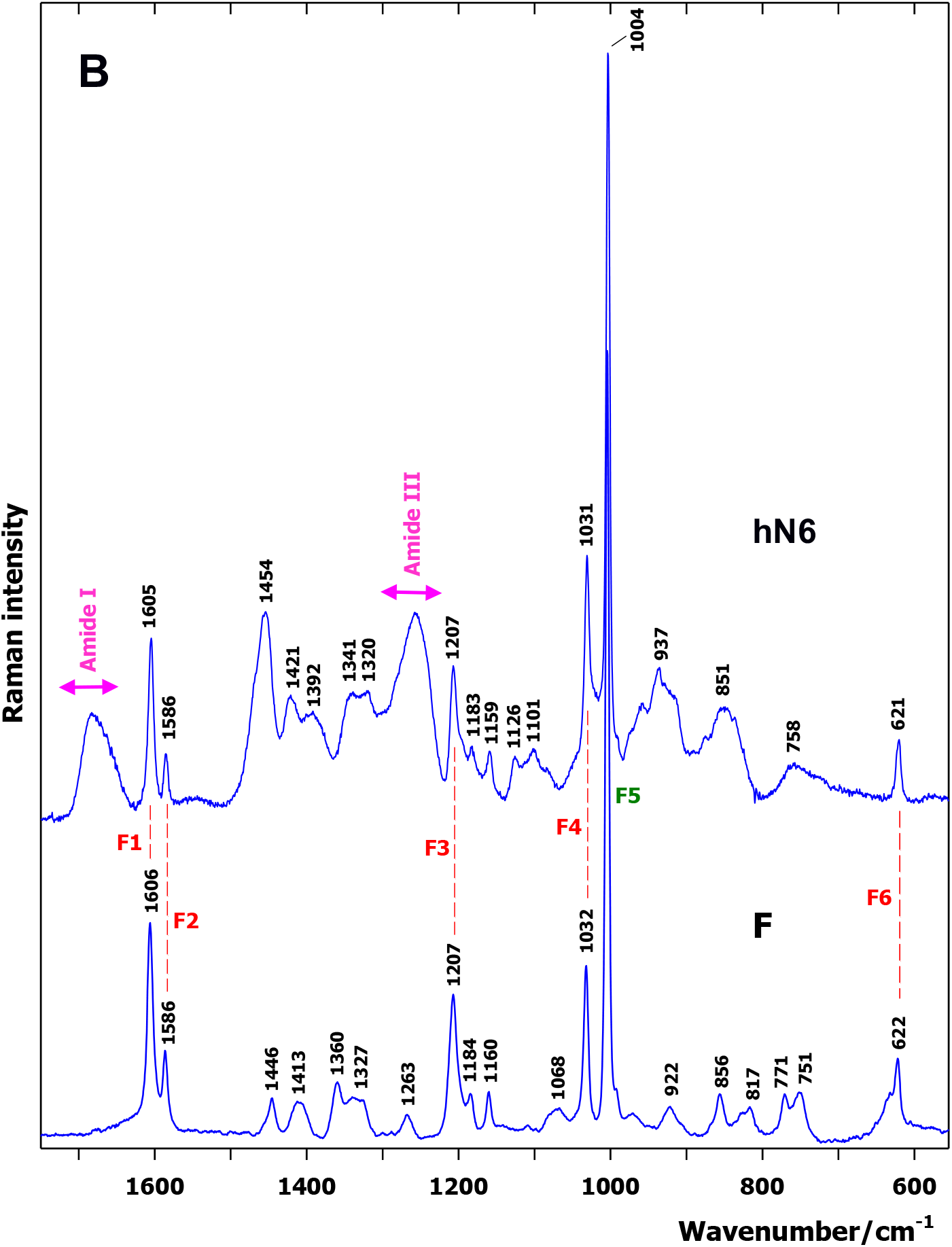
Room temperature Stokes Raman spectra in the middle wavenumber region for (A) hN3, hN4, hN5 and (B) hN6, compared to F. Spectra were recorded at 20 mM. See Table 2 for tentative assignments.

The tentative assignments of the Raman spectra of the four studied nonapeptides are reported in Table 2. These assignments have been performed in keeping with the features observed in the Raman spectra of their constitutive amino-acids (Fig. S1). The bands observed in the 1500-1320 cm^−1^ range, mainly arising from the bending modes of CH_2_ and CH_3_ side chain moieties, as well as those located below 1200 cm^−1^ in amino acids, were taken as references for assigning the observed peaks in the spectra of peptides.

**Table 2.** Tentative assignment of the Raman bands observed in the middle wavenumber region.

| **AGIPGLGVG**<br>**hN3** | **VGVPGLGVG**<br>**hN4** | **AGVPGLGVG**<br>**hN5** | **AGVPGFGAG**<br>**hN6** | **Tentative**<br>**assignment** |
| --- | --- | --- | --- | --- |
| 1700-1620 (br) | 1700-1620 (br) | 1700-1620 (br) | 1700-1620 (br) | Amide I |
|  |  |  | 1605 (s) | F1 |
|  |  |  | 1586 (m) | F2 |
| 1465 (s) | 1468 (s) | 1466 (s) |  | P, I, L, V, A |
| 1451 (s) | 1452 (s) | 1452 (s) | 1454 (s) | V, L, I |
| 1421 (s) | 1421 (s) | 1420 (s) | 1421 (s) | G, A, L, I, V, P |
| 1391 (m) | 1394 (m) | 1395 (m) | 1392 (m) | G, A, L, I, V, P |
| 1343 (s) | 1345 (s) | 1343 (s) | 1341 (s) | A, L, I, V, P |
| 1315 (s) | 1313 (s) | 1317 (s) | 1320 (s) | A, L, I |
| 1300-1240 (br) | 1300-1240 (br) | 1300-1240 (br) | 1300-1240 (br) | Amide III |
|  |  |  | 1207 (s) | F3 |
|  |  |  | 1183 (m) |  |
| 1161 (m) | 1175 (m) | 1161 (m) | 1159 (m) | V, L, I, P |
| 1128 (s) | 1127 (s) | 1126 (s) | 1126 (m) | V, L, I |
|  |  |  | 1101 (m) | P |
|  |  |  | 1031 (s) | F4 |
| 1031 (m) | 1034 (m) | 1031 (m) |  | G, I, P |
| 1011 (m) | 1014 (m) | 1014 (m) |  | A, V |
|  |  |  | 1004 (vs) | F5 |
| 960 (s) | 962 (s) | 960 (s) |  | V, L, I |
| 937 (s) | 938 (s) | 937 (s) | 937 (br) | A, V, L, I |
| 891 (s) | 891 (s) | 891 (s) |  | V |
| 857 (m) | 860 (m) | 856 (sh) | 851 (br) | L, P |
| 845 (m) | 840 (m) | 843 (m) |  | A, V, L, I, P |
|  |  |  | 758 (br) | P |
| 736 (br) | 735 (br) | 737 (br) |  | V, L, I, P |
|  |  |  | 621 (m) | F6 |

(vs) very strong; (s) strong; (m) medium; (br) broad Raman bands.
F1 to F6 refer to the six characteristic Raman bands of phenylalanine. Amide I and amide III vibrations give rise to a broad and strong Raman band peaking at $\sim 1685$ and $1255\text{ cm}^{-1}$ , respectively.

The spectrum of hN6 is characterized by the occurrence of the six Phe characteristic Raman bands, referred to as F1 to F6 (40, 41). Being all of in-plane type and localized in the Phe phenyl ring, the wavenumbers of these six characteristic markers remained very close to those collected from the free amino acid F (Fig. 3B). The Raman spectra of hN3, hN4 and hN5 (Fig. 3A) do not present such narrow and resolved aromatic markers, but their striking spectral shape similarity is to be emphasized (Fig. 3). Two regions corresponding to amide I (1700-1640 cm^−1^) and amide III (1300-1230 cm^−1^) vibrations provide valuable structural information through their decomposition into band components (36–38). It is noteworthy that while amide I vibrations results from the backbone C=O bond stretch motion, more or less coupled to its adjacent N-H angular bending, amide III vibrations mainly arise from the backbone N-H angular bending. The Raman spectra obtained for the free amino acids (Fig. S1) are devoid of amide (I and III) vibrations. However, the presence of low intensity bands arising from the side chains of Val, Leu, Ile and Pro, falling within the amide III range should be stressed. These bands can be naturally superimposed to those relative to amide III vibrations in peptides. Nevertheless, because of their weakness, one cannot expect a considerable distortion in the structural analysis on the basis of amide III vibrations. In all nonapeptides, amide I and amide III vibrations are characterized by two strong, broad and incompletely resolved bands peaking at ~1685 and ~1265 cm^−1^, respectively. To go farther in extracting the structural information from amide vibrations, the decomposition of the amide I region for four nonapeptides is displayed in Figure 4. Through a systematic investigation on the structural analysis of the peptide chains (36–38, 42, 43), in aqueous environment, we have been led to select four band components located at ~1690±5 (assignable to random chain), ~1660±5 (assignable to β-strand), ~1675±5 and ~1650±5 (both assignable to turns) to decompose the amide I region in nonapeptides. The populations corresponding to different secondary structural elements, as estimated by the normalized areas of the band components (expressed in percent) are reported in Table 3. As it can be seen between 50% to 60% of the total population is formed by turn structures, and the rest is equally distributed between β-strands and random chains.

**Table 3.** Underlying secondary structure elements in the Amide I profile of the nonapeptides and fractional areas.

| Position (assignment) | AGIPGLGVG | VGVPGLGVG | AGVPGLGVG | AGVPGFGAG |
| --- | --- | --- | --- | --- |
| ~1690 (random) | 20 | 20 | 24 | 25 |
| ~1680 (turn) | 41 | 40 | 37 | 38 |
| ~1660 (β-strand) | 26 | 23 | 21 | 25 |
| ~1640 (turn) | 13 | 17 | 18 | 12 |
| Fractional areas |  |  |  |  |
| Random | 20 | 20 | 24 | 25 |
| Turn | 54 | 57 | 55 | 50 |
| β-strand | 26 | 23 | 21 | 25 |

Component widths were kept between 15 and 25 $\text{cm}^{-1}$ . The figures correspond to the normalized areas of the components expressed in per cent of the whole Amide I profile. Fractional areas estimate the global secondary structures of the peptide.

**Figure 4.**
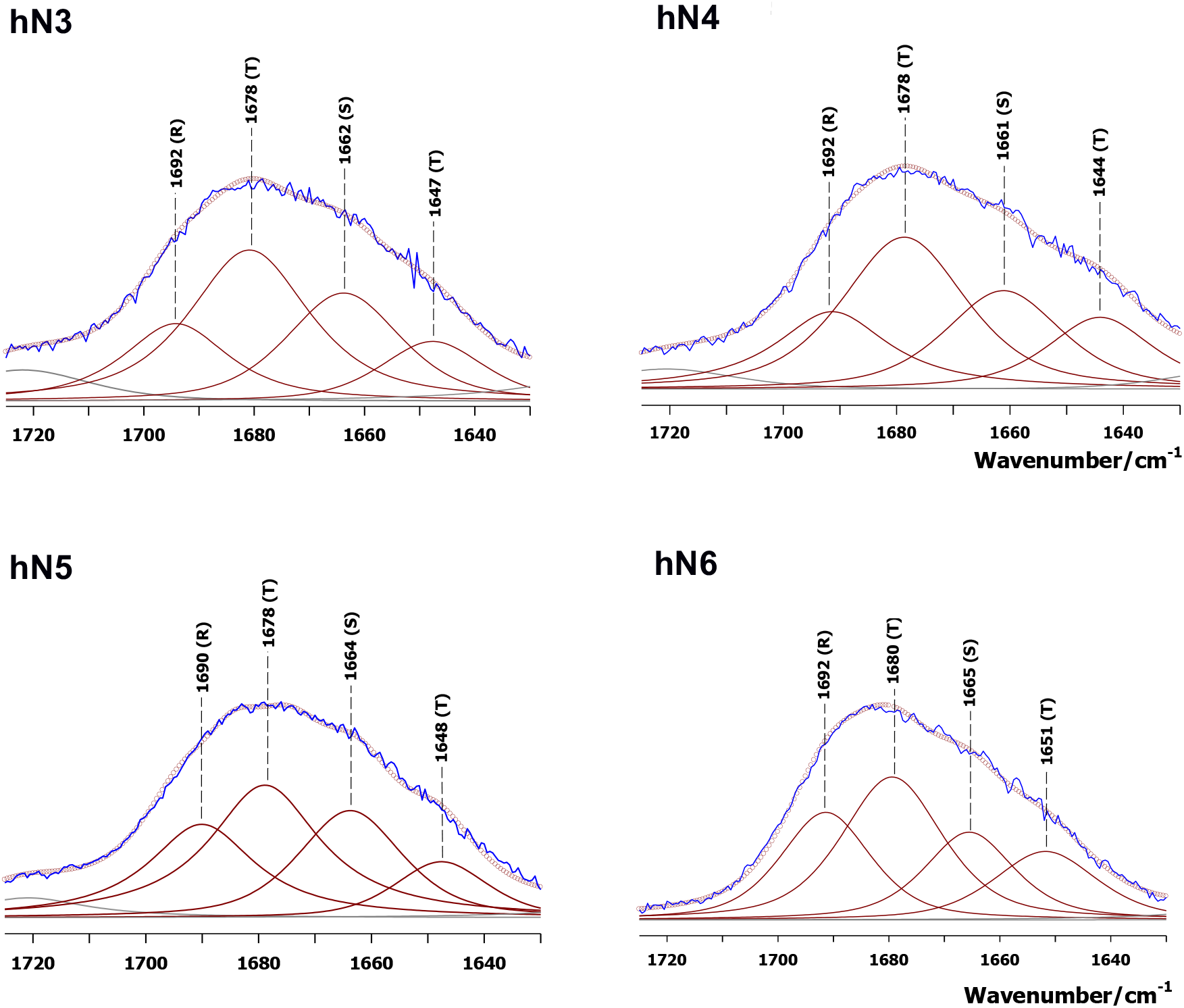
Band decomposition in the amide I region of the nonapeptides observed at 20 mM. Observed spectra are drawn with blue lines, component bands with red lines, red circles correspond to the sum of component bands. Maximum band wavenumber of each used component is reported. In parentheses: (R) random; (T) turn; (S) β-strand. For the estimated populations of different secondary structural elements, see Table 3.

### Structural behavior as a function of temperature

Solution samples containing nonapeptides at 20 mM were heated up to 80°C. Little changes appeared in their Raman spectra, but slight changes could be observed in the amide III region. Figure 5 exemplifies this behavior for hN5 and hN6. The main effect is a wavenumber downshift of the spectral profile consistent with a progressive shift of the conformational equilibrium from turn to β-strand upon increasing temperature. Albeit the turns remain the major population of secondary structures in the 10-80° C temperature range (data not shown), they show a little tendency to be transformed into β-strands, presumably because of the weakening/breakdown of the inter-turn stabilizing hydrogen bonds occurring upon thermal annealing.

**Figure 5.**
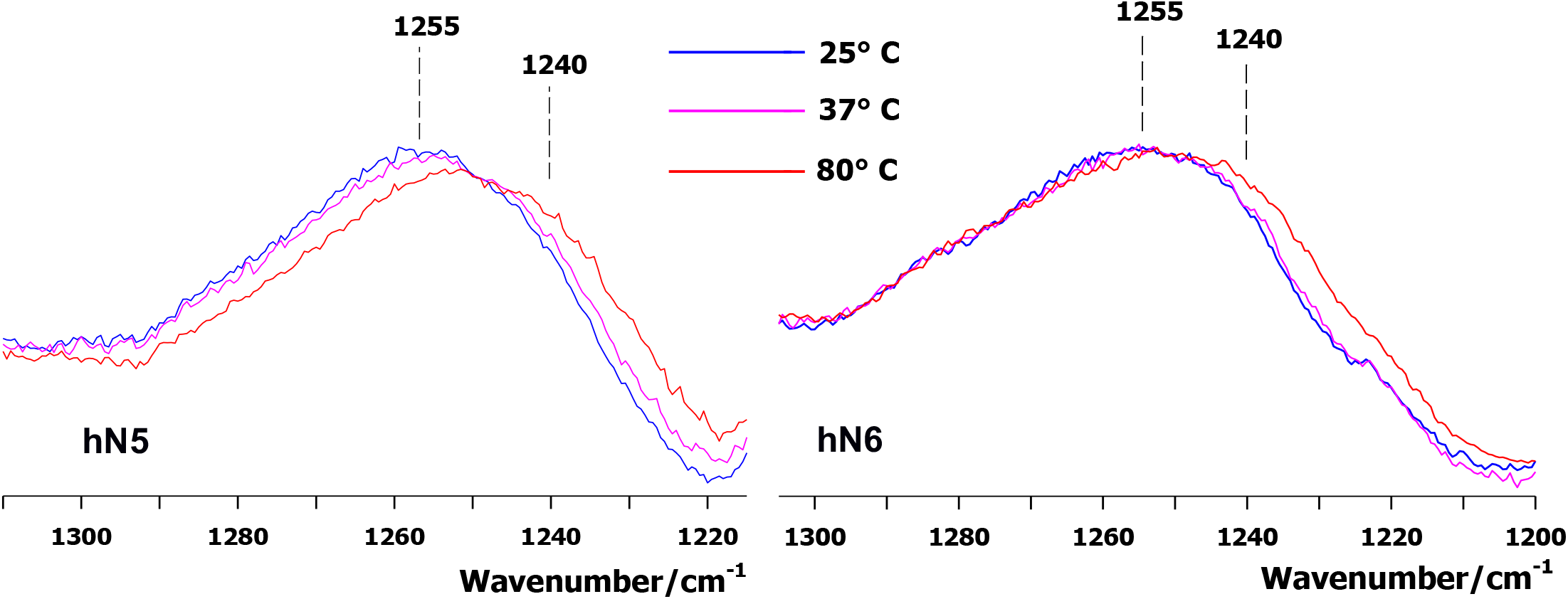
Effect of temperature on the amide III vibrations of nonapeptides. The spectra were recorded for a concentration of 20 mM. Different colors are used to display the spectra obtained as a function of temperature.

### Atomistic molecular dynamics simulations

Molecular dynamic simulations were performed for the four nonapeptides. Figure 6A presents evolution of the conformation computed on each position of the nonapeptide hN5 throughout the 500 ns duration of the simulation. Similar results for hN3, hN4 and hN6 are provided as Figure S2. This figure shows that no stable structure can be observed in the peptide. The peptide adopts mainly random coil conformations with residues favoring β-turns mainly in the middle of the peptide, notably at positions 4 (P) and 5 (G). As the extremities of the peptides are mainly random coil, distances have been computed between C_α_ of residues 3 and 7. Analysis of the simulation underlines that the structures are rapidly changing on the nanosecond time scale from extended to more packed conformations. However, the distance distribution (Fig. 6B) shows, for each nonapeptide, the existence of a population of conformations with a C_α3_-C_α7_ distance of about 0.5 nm. This well-defined population corresponds to the transient formation of β-turns. Indeed, structure clustering (Fig. 6C) shows that this population adopts a β-turn. It is the dominant structural population for hN3 (48.5 ± 3.7%), hN4 (35.1 ± 4.3%), hN5 (31.8 ± 3.6%) and hN6 (27.5 ± 3.1%). In our simulations, the formation of turns is also associated with an increase of the number of local hydrogen bonds. The greater number is observed for hN3 and in a lesser extend for hN4 and hN5. Altogether, our analysis of MD data strongly suggests that the PG motif of the nonapeptides could mainly adopt a β-turn conformation.

**Figure 6.**
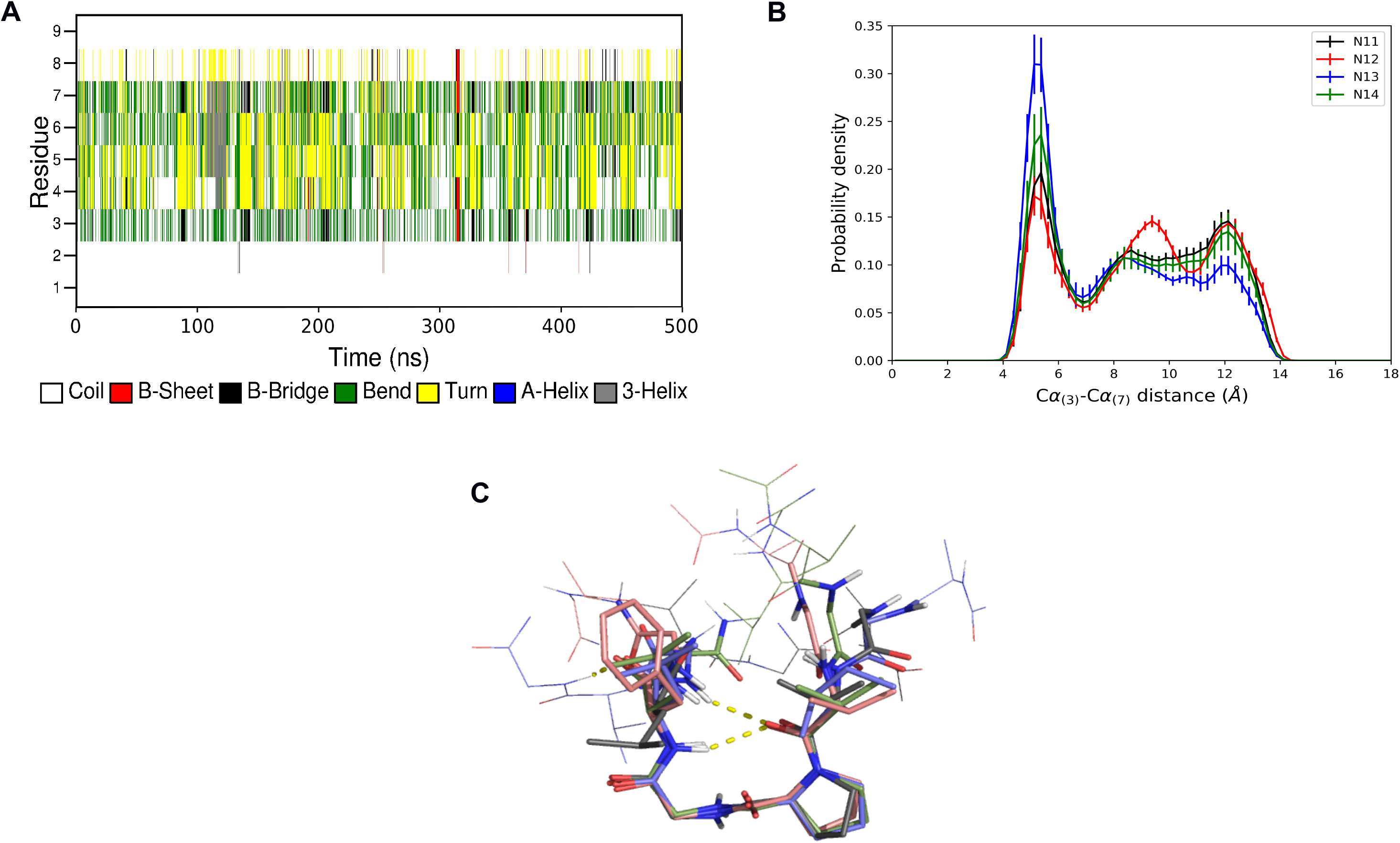
Conformational analysis following molecular dynamics simulations. A, secondary structures adopted during the simulation by hN5. B, distribution of distances between the third and seventh residues for the four nonapeptides. C, overlay of the conformation representative of the main cluster computed for the four nonapeptides. Residues 3 to 7 are represented with a thicker line.

## Discussion

Among elastin peptides, elastin nonapeptides have been identified initially as chemo-attractants for fibroblasts (12). Now, there are growing evidences that these elastin peptides could define a peculiar class of matrikines.

Brassart and colleagues have recently shown that hN5 (*i.e.* AGVPGLGVG) plays a key role in tumor progression by promoting tumor cell blebbing and extracellular vesicle shedding following its interaction with the ribosomal protein SA (17). The structure of hN5 has been analyzed previously by CD, FT-IR and NMR spectroscopies (16). This report concluded that the structure of the peptide was a mixture of random coils and β-turns. Our data fully agree with these findings but they also expand our knowledge of the nature and structural behavior of these elastin peptides.

The occurrence of the X-G-X-P-G-X-G-X-G consensus sequence was analyzed in a large sequence data bank. Our results show that among the observed hits, the group of elastin sequences defines a specific group (Fig. 1). Further, our analysis underlines the fact that these peptides mostly occur in tandem repeats in human exon 26 (and corresponding sequences in other species). Finally, as far as human sequences are concerned, we observed that similar sequences were also present in numerous species (Table 1). The most striking observation is that hN6 is present in numerous species and is tandemly repeated five times in mouse sequence. In contrast, hN3 is only found in 3 species. These observations suggest that, besides their matrikines activity, these sequences could play a key, and yet undescribed, role in elastin biology, possibly elastin assembly. Indeed, among elastin sequences, these sequences appear much more conserved that the canonical elastin hexapeptide sequence (say VGVAPG), which occurrence is mostly limited to human exon 24 sequence. Further works are needed to test this hypothesis.

In the present study, we focused our analysis on the nonapeptidic sequences found in human exon 26 aiming at understanding their structure and structural behavior. First, this investigation was performed using optical spectroscopies. The results gathered by both CD and Raman spectra is consistent with previous reports underlining conformational equilibrium existing between random, sheet and turns conformations within elastin (44). Nevertheless, the Raman data underline that the conformation of these peptides is dominated by β-turns as deduced from the quantitative analysis of their amide I profiles. These conformations appear very stable as increasing the temperature up to 80°C did not drastically change the spectra and hence their structure, albeit a small increase of the β-sheets content could be observed upon heating. Altogether our experimental data suggest that the structures of the nonapeptides are dominated by β-turns and that these conformations are stable even though engaged in a conformational equilibrium with extended and random structures.

In order to have a better understanding of the structural equilibrium occurring in these peptides, molecular dynamics simulations were undertaken in explicit water. Ten 500-ns simulations were performed for the 4 nonapeptides. The results obtained are extremely consistent with our experimental data. Indeed, they show that the most common structures observed during the simulation is a β-turn occurring at the X-P-G-X motif. Quantitatively, this structure seems more abundant in hN3 (about 50%), than hN4, hN5 or hN6 (25-35%). This finding is in very good agreement with the conclusions of Lessing and colleagues (45) who observed that the preference for the X-Pro-Gly sequence to form a β-turn increased with the complexity and hydrophobicity of the side chain at position X (Gly < Ala < Val). In contrast to hN4, hN5 and hN6 which harbor a valyl residue before PG, hN3 possesses an isoleucyl residue at that position. Thus, the higher propensity of hN3 to adopt a β-turn when compared to the other three nonapeptides is consistent with the work of Lessing and colleagues.

Since the first report of their bioactivity on bovine fibroblasts (12), nonapeptides sequences have been tested on various cellular systems: bovine endothelial cells (13), rat macrophages (14) and human tumor cell lines (8, 9). Nevertheless, among those publications few used more than one peptide sequence allowing comparison of their respective effects. In the work of Maeda and colleagues, hN5 and hN6 were identified as chemoattractant for rat macrophages (14). In this work, the author also analyzed the effect of the canonical VGVAPG sequence. Their conclusion was that the two nonapeptides could compete for receptor binding. It has been established that VGVAPG bioactivity relies on its ability to adopt mainly a type VIII β-turn conformation (10). Our molecular dynamics simulations show that their major conformation could be the type II β-turn. This apparent contraction with our theoretical calculations is however balanced by our experimental Raman data as they suggest that type VIII β-turn could also be present in the conformation of the nonapeptides.

Altogether our data suggest that the structural signature of elastin nonapeptide is a type II β-turn. Consequently, it is reasonable to propose that this conformation could be relevant of the observed bioactivity of these peptides. This proposal is supported by the observations made by Toupance and co-workers where they measured a lesser biological effect of hN6 towards human tumor cells than hN5 (8). In our work, hN6 is predicted to present less β-turn conformations than hN5. Therefore, it seems coherent that the bioactivity of hN5 is greater than that of hN6 supporting the proposal that the turn motif is the conformation sustaining bioactivity of these peptides. If our prediction is correct, then one can anticipate that hN3 would be a peptide more bioactive that others. Further work is need to test this hypothesis.

### Conclusion

Our data throw new light on the nature, structure and conformational behavior of elastin nonapeptides. These peptide sequences exhibit a peculiar composition as demonstrated by our bioinformatic analysis of current databases. Further, experimental (CD and Raman spectroscopies) and theoretical data (molecular dynamics) strongly suggest that the dynamic structure of these peptides is dominated by type II β-turn. In keeping, we propose that the observed bioactivity of these peptides towards stromal and tumor cells could be driven by the recognition of this structural feature by their cognate receptor, the ribosomal protein SA. Molecular modeling of their interaction is underway.

## Authors contributions

HB and KSG performed the Raman and CD measurements and analyzed the spectra. BS conceived and ran the simulations. CJM, TJ and BN performed the sequence analyses. BS and DM funded the project and edited the manuscript. DL supervised the work and wrote the manuscript.

## Acknowledgments

The authors would like to thank the MAgICS chair (Université de Reims Champagne Ardenne) and the Centre National de la Recherche Scientifique (CNRS) for financial support. Molecular dynamics calculations were performed on the ROMEO high performance computing center.

